# The phosphorylation status of LRRK2 at the S910/S935 cluster determines its sensitivity to activation by RAB29

**DOI:** 10.64898/2026.05.22.727151

**Authors:** Antonio Jesús Lara Ordóñez, Chloé Annicotte, Enna Behrends, Margaux Morez, Alessio Burin, Liesel Goveas, Femke Van Mele, Christian Galicia, Wim Versées, Jean-Marc Taymans

## Abstract

Leucine-Rich Repeat Kinase 2 (LRRK2) is a signaling molecule involved in Parkinson’s disease pathomechanisms. In disease, the LRRK2 protein displays both a toxic gain of kinase function and a loss of phosphorylation at heterophosphosites found in an extended loop of the LRR domain. RAB GTPases, such as RAB29, have been identified as upstream activators of LRRK2. Indeed, co-expression of LRRK2 with RAB29 induces a hyperactivation of LRRK2 kinase activity, however the role of the LRRK2 heterologous phosphorylation status in its activation remains unknown. Here, our aim was to determine the role of LRRK2 heterologous phosphorylation on its activation by RAB29. Using single and compound phosphodead or phosphomimetic mutants of LRRK2 we show differential sensitivity of LRRK2 phosphomutants to activation by RAB29, with phosphodead mutants being more susceptible to be activated than phosphomimetic mutants. Interestingly, we find that the single phosphodead S910A LRRK2 mutant displays an activation of LRRK2 kinase activity similar to that observed for the compound phosphodead 6xS>A LRRK2 mutant (S860A/S910A/S935A/S955A/S973A/S976A). Time-course analysis revealed that phosphodead mutants displayed higher but also faster activation by RAB29. In addition, both physical interaction between LRRK2 and RAB29 as well as RAB29-induced recruitment of LRRK2 to the *trans*-Golgi network (TGN) was enhanced by phosphodead compared to phosphomimetic mutants. To confirm effects on native LRRK2, we tested a panel of ten nanobodies targeting LRRK2 that stabilized LRRK2 phosphorylation at varying levels. Nanobodies stabilizing LRRK2 at low S935 phosphorylation levels showed enhanced RAB29-induced activation compared to nanobodies not affecting pS935 LRRK2. Finally, we tested whether LRRK2 heterologous phosphorylation could affect centrosome cohesion deficits, a phenotype that has been linked to LRRK2 hyperactivation, and found that both the phosphodead LRRK2 as well as a nanobody stabilizing dephosphorylated LRRK2 enhanced the centrosome cohesion deficit. Our findings indicate that hyperactivability of LRRK2 is directly related to its heterologous phosphorylation status, with dephosphorylation leading to strong hyperactivation of LRRK2 by upstream activating RABs, and phosphorylated LRRK2 showing the opposite. This implies that strategies favoring LRRK2 phosphorylation will have therapeutic benefit.

## Introduction

Leucine-rich repeat kinase 2 (LRRK2) is a signaling molecule involved in intracellular organization that has been implicated in Parkinson’s disease (PD), Crohn’s disease, leprosy and cancer (Wang et al., 2026). In PD, evidence from biochemical, cellular and animal models as well as the analysis of patient samples show that pathology is correlated to the hyperactivation of LRRK2. Among other evidences, relatively rare LRRK2 clinical mutants demonstrate an enhanced kinase activity compared to WT LRRK2 (Steger et al., 2016) and LRRK2 kinase activation has also been observed in *post-mortem* brains of PD patients (Di Maio et al., 2018). LRRK2 has been considered as a priority therapeutic target for Parkinson’s disease, and therapeutic strategies being developed are based on the inhibition of LRRK2 kinase activity (Morez et al., 2024). Indeed, LRRK2 kinase inhibitors have been developed and are currently in clinical trials (Morez et al., 2024; Lang et al., 2026). Recently, interest has grown to explore alternative targeting methods to block LRRK2 hyperactivation, however information on activation mechanisms of LRRK2 is fragmentary.

Early studies of enzymatic activity of clinical pathogenic mutants of LRRK2 found that these have enhanced kinase activity compared to WT, under basal conditions (Greggio and Cookson, 2009). While enhanced basal kinase activity is well established as a disease mechanism, it is clear that additional layers of regulation may exist, for instance to explain the role of native LRRK2 in PD or to explain the lack of full penetrance of disease mutants. In recent years, some specific conditions have been found to lead to LRRK2 activation. Treatment with lysosomotropic agents such as chloroquine or L-leucyl-L-leucine methyl ester (LLOMe) lead to an increase of phosphorylation of LRRK2’s substrate RAB10 at T73, considered a cellular marker of LRRK2 kinase activity (Kuwahara et al., 2020). In addition, several upstream activators of LRRK2 have been identified, including RAB12, RAB29 and VPS35, which when overexpressed lead to upregulation of markers of activated LRRK2 kinase, including both LRRK2 autophosphorylation at S1292 and RAB phosphorylation (Mir et al., 2018; Purlyte et al., 2018; Dhekne et al., 2023). Interestingly, RAB-mediated activation is enhanced to varying degrees in disease mutants of LRRK2 compared to WT, suggesting that specific mutants of LRRK2 are more susceptible to activation than others (Kalogeropulou et al., 2022).

In the present study, we address the question of how activation of LRRK2 is affected by LRRK2’s differential phosphorylation. LRRK2 is phosphorylated on multiple sites, either through autophosphorylation or through phosphorylation by heterologous kinases (Marchand et al., 2020). Heterologous phosphorylation sites cluster in an extended loop between leucine-rich repeat segments of the LRR domain (heterophospholoop). This cluster includes several phosphoserines, including at sites S910 and S935, that coordinate binding to 14-3-3 proteins. These sites are intimately linked to LRRK2 function and disease, as pS910/pS935 levels are reduced in several clinical mutant forms of LRRK2, in *post-mortem* PD brain as well as *in cellulo* and *in vivo* models after treatment with LRRK2 kinase inhibitors (Dzamko et al., 2010, 2017; Vancraenenbroeck et al., 2014; Di Maio et al., 2018; Marchand et al., 2020). One hypothesis that has previously been addressed is that LRRK2 heterophosphorylation may be correlated to LRRK2 kinase activity in basal conditions. This was tested on several clinical and functional mutants of LRRK2 with varying effects on kinase activity and pS935 LRRK2 levels, with results revealing a lack of correlation between pS935 LRRK2 and activity in basal conditions (Ito et al., 2014). Previous work to address this question has also been done using phosphomutant forms of LRRK2, replacing the serine by an alanine to mimic dephosphorylation (phosphodead) or by an aspartate to mimic phosphorylation (phosphomimetic). Using compound hexamutants of the phosphorylation loop of the LRR (S860/S910/S935/S955/S973/S976), we recently found that phosphomimetic 6xS>D LRRK2 displayed reduced kinase activation in basal conditions, and that phosphodead 6xS>A LRRK2 showed an increased propensity for activation by the LRRK2 activator RAB29 (Marchand et al., 2022).

Therefore, our aim in this study is to dissect out the role of individual phosphosites of the heterophospholoop of LRRK2 during LRRK2 hyperactivation and how this affects downstream molecular and cellular phenotypes. Since it is not clear whether individual sites within this loop are responsible for altered sensitivity of LRRK2 to activation or whether the whole loop is involved, we employed a phosphomutant approach to mimic LRRK2 phosphorylation or dephosphorylation at specific sites. In addition, to confirm links between LRRK2 phosphorylation and activation on native LRRK2, we used a panel of LRRK2-targeting nanobodies (Nbs) (Singh et al., 2022) that are known to stabilize proteins in specific conformations and that we found to have differential effects on LRRK2 phosphorylation. We subjected both LRRK2 phosphomutants and LRRK2 with phosphoregulating nanobodies to activation conditions by upstream activator RAB29, and investigated how this is linked to kinase activity markers and other downstream cellular effects such as LRRK2 recruitment to organelles and LRRK2-mediated centrosome cohesion deficits.

## Materials and methods

### DNA constructs

Empty (CMV) and 3xFlag-tagged human LRRK2 WT and phosphomutant lentiviral constructs have been previously described (Marchand et al., 2022). Nanobody constructs for eukaryotic expression of Nbs with tandem 6xHis-eGFP tag have been previously described (Singh et al., 2022). peGFP-C1 was obtained from Clontech. eGFP-tagged RAB29 was kindly provided by Dr Sabine Hilfiker (Rutgers New Jersey Medical School, NJ, USA) (Madero-Pérez et al., 2018b). All constructs were prepared from bacterial cultures grown at 37°C using PureYield™ Plasmid Midiprep System (Promega, A2495) according to manufacturer’s instructions. Plasmids were verified by sequencing of the entire coding region (Eurofins Genomics GmbH, Germany).

### Cell culture and transfection

HEK293T cells (ATCC, CRL-3216) were cultured as previously described (Marchand et al., 2022). Briefly, cells were grown in Dulbecco’s modified Eagle’s medium containing high glucose (Sigma-Aldrich, D6429), supplemented with 10% fetal bovine serum (Fisher Scientific 15595309), non-essential amino acids (Fisher Scientific, 12084947), and 100 U/ml penicillin and 100 μg/ml streptomycin (Fisher Scientific, 15140122). Cells were subcultured in 12-well plates and transfected at 70-80% confluence with 1 µg of LRRK2 constructs and 100 ng of eGFP or eGFP-RAB29 (and 500 ng of Nbs where indicated) using polyethylenimine (PEI, branched) (Sigma-Aldrich, 408727). The next day, cells were collected or split onto poly-D-lysine coated coverslips (Sigma-Aldrich, P0899) and subjected to immunocytochemistry or western blot analysis 24 h or 48 h after transfection. For co-immunoprecipitation, cells were seeded in 100mm dish, transfected at 70-80% confluence with 12 µg of LRRK2 constructs (and 2 µg of eGFP or eGFP-RAB29 where indicated) using polyethylenimine (PEI, branched) (Sigma-Aldrich, 408727), and subjected to immunoprecipitation and western blot analysis 24 h or 48 h after transfection.

Where indicated, cells were treated with DMSO (Sigma-Aldrich, D2438) or MLi-2 (Tocris, 5756).

### Immunocytochemistry

Cells were fixed with 4% paraformaldehyde (PFA) in PBS for 15 minutes at room temperature, and permeabilized with 0.2% TritonX-100/PBS for 10 minutes at room temperature, followed by incubation with blocking solution (0.5% BSA (w/v) in 0.2% TritonX-100/PBS) for 1 hour at room temperature. Coverslips were incubated with the indicated primary antibodies diluted in blocking solution overnight at 4°C. Primary antibodies included mouse monoclonal anti-Flag® M2 (1:500, Sigma-Aldrich, F1804), sheep anti-human TGN46 (1:2000, Bio-Rad, AHP500G), and rabbit polyclonal anti-pericentrin (1:1000, Abcam, ab4448). The following day, coverslips were washed twice with 0.2% TritonX-100/PBS for 10 minutes at room temperature and incubated with the indicated secondary antibodies diluted in 0.2% TritonX-100/PBS for 1 hour at room temperature. Secondary antibodies included Alexa Fluor™ 488 goat anti-mouse (1:1000, Invitrogen, A11001), Alexa Fluor™ 568 donkey anti-mouse or goat anti-rabbit (1:1000, Invitrogen, A10037 or A11011), and Alexa Fluor™ 647 donkey anti-sheep (1:1000, Invitrogen, A21448). After two additional washes, coverslips were rinsed in distilled water and 70% ethanol, air-dried and mounted with VECTASHIELD® Antifade Mounting Medium with DAPI (Vector Laboratories, H-1200).

### Laser confocal imaging and analysis

For LRRK2 localization to TGN, images were acquired on a Zeiss LSM710 confocal microscope using a 63x 1.4 NA oil-immersion objective (HCX PLAPO CS). Images were collected using single excitation for each wavelength separately and dependent on the secondary antibodies (561 nm DPSS Laser line and a 610-640 nm emission band pass, and 633 HeNe Laser line and a 655-710 nm emission band pass), eGFP-tagged proteins were excited with 488 nm Argon Laser line (and a 500-540 nm emission band pass), and DAPI was excited with 405 nm UV Diode (and a 420-465 nm emission band pass). To optimize signal detection, laser intensity settings and exposure time were adjusted for each channel according to the emitted fluorescence intensity.

Twenty to twenty-five image sections of selected areas were acquired with a step size of 0.5 µm, and Z-stack images analyzed and processed using Fiji software (Schindelin et al., 2012). Quantification was based on the number of individual cells that displayed co-localization of LRRK2, RAB29 and TGN46, relative to the total number of cells co-transfected with LRRK2 and RAB29 in the indicated condition. At least 60 co-transfected cells with LRRK2 and RAB29 were analyzed per condition per experiment blind to condition.

For centrosome cohesion, images were acquired on a Leica TCS SP8 MP confocal microscope using a 63x 1.4 NA oil-immersion objective (HCX PLAPO CS). Images were collected using single excitation for each wavelength separately and dependent on the secondary antibodies (552 nm OPSL Laser line and a 590-620 nm emission band pass, and 638 nm Diode and a 655-685 nm emission band pass), eGFP-tagged proteins were excited with 488 nm OPSL Laser line (and a 500-530 nm emission band pass), and DAPI was excited with 405 nm UV Diode (and a 420-470 nm emission band pass).

Twenty to twenty-five image sections of selected areas were acquired with a step size of 0.5 µm, and Z-stack images analyzed and processed using Leica Applied Systems (LAS X Office) image analysis software. Centrosomes were scored as being separated when the distance between their centers was >1.5 μm (Madero-Pérez et al., 2018a; Lara Ordóñez et al., 2019). In all cases, mitotic cells were excluded from the analysis.

### Cell lysis, co-immunoprecipitation and western blot

Cells were collected and washed in PBS, followed by resuspension in ice-cold lysis buffer (20 mM Tris pH 7.5, 150 mM NaCl, 0.2% Triton-X100, 5 mM MgCl_2_, 10% glycerol) supplemented with protease and phosphatase inhibitors (cOmplete™ EDTA-free Protease Inhibitor Cocktail and PhosSTOP™, Sigma-Aldrich, 046931320019 and 4906837001, respectively) and incubation on rotary wheel for 30 minutes at 4°C. Lysates were then centrifuged at 13,200 rpm for 10 minutes at 4°C, and protein concentration of supernatants estimated by BCA assay (Takara, T9300A) according to manufacturer’s instructions. For co-immunoprecipitation assay, a small sample of lysate was set aside to determine input expression levels, while the rest of the lysate was incubated with 15 µL of ChromoTek GFP-Trap® Magnetic Agarose (Proteintech, gtma-20) for 30 minutes at 4 °C on a rotary wheel. After incubation, samples were washed 3 times with lysis buffer. 30 µL of a solution of 2X NuPAGE™ LDS Sample Buffer and Bolt™ Sample Reducing Agent (Invitrogen, NP0008 and B0009, respectively) were added to each sample, followed by vortexing, heat treatment at 95 °C for 10 minutes and final vortexing. These steps allowed to detach the proteins of interest, alongside the antibodies used for capture, from the magnetic beads. Samples were then analyzed by western blot.

Cell lysates were mixed with the appropriate amount of a solution of 2.85X NuPAGE™ LDS Sample Buffer and Bolt™ Sample Reducing Agent (Invitrogen, NP0008 and B0009, respectively) and boiled at 95°C for 5 minutes (or 70°C for 10 minutes). Around 25 µg of protein were resolved in NuPAGE™ 4-12% Bis-Tris Mini or Midi Gels (Invitrogen, NP0329 and WG1403, respectively) at 100V for 2 hours with NuPAGE™ MOPS SDS running buffer (Invitrogen, NP0001), and electrophoretically transferred onto 0.45 µm nitrocellulose membranes (Cytiva, 10600002) at 40 mA for Mini gels or 120 mA for Midi gels overnight at 4°C in transfer buffer (20 mM Tris pH 8.6, 122 mM glycine, 5% (v/v) MeOH). Membranes were blocked with 5% (w/v) BSA or 5% (w/v) skim milk powder in 0.1% Tween-20/PBS for 1 hour at room temperature, and incubated with the indicated primary antibodies diluted in blocking solution overnight at 4°C. Primary antibodies included rabbit monoclonal anti-pS910 LRRK2 [UDD1 15(3)] (1:1000, Abcam, ab133449), rabbit monoclonal anti-pS935 LRRK2 [UDD2 10(12)] (1:1000, Abcam, ab133450), rabbit monoclonal anti-pS1292 LRRK2 [MJFR-19-7-8] (1:1000, Abcam, ab203181), mouse monoclonal anti-Flag® M2 (1:500, Sigma-Aldrich, F1804), mouse monoclonal anti-α-tubulin (1:10000, Bio-Techne, NB100-690), rabbit polyclonal anti-GAPDH (1:5000, Sigma-Aldrich, G9545), rabbit polyclonal anti-GFP (1:1000, Invitrogen, A11122), rabbit monoclonal anti-pT73 RAB10 [MJF-R21] (1:1000, Abcam, ab230261), and mouse monoclonal anti-RAB10 [4E2] (1:1000, Invitrogen, MA5-15670). The following day, membranes were washed twice in 0.1% Tween-20/PBS for 10 minutes at room temperature and incubated with secondary antibodies diluted in 0.1% Tween-20/PBS for 1 hour at room temperature. Secondary antibodies included HRP-conjugated goat anti-mouse (1:10000, Cell Signaling, 7076S) and goat anti-rabbit (1:5000, Cell Signaling, 7074S). After two additional washes, membranes were revealed using ECL Standard or Prime Western Blotting Detection Reagent (Cytiva, RPN2209 or RPN2236, respectively). Protein bands were detected with either Amersham ImageQuant™ 600 or 800 western blot imaging systems (Cytiva, USA), and analyzed using either Amersham ImageQuant™ software version 10.2 (Cytiva, USA) or Image Studio™ Lite software version 5.2 (LI-COR Biotech, USA).

### Structural modelling

AlphaFold 3 (AF3) (Abramson et al., 2024) and Chai-1 (Chai Discovery, 2024) were used to model binding positions of Nb23 to LRRK2. AF3 was used with default inference parameters, while the sequence of either full-length LRRK2 or only its kinase-WD40 domains was used as input. Chai-1 was run using the kinase-WD40 domains and using constrained-based modelling in its two modalities, “pocket-mode” and “binding-mode”, both yielding similar results. In the binding-mode, constraints were defined based on the results of previous crosslinking mass spectrometry experiments (Singh et al., 2022), setting a maximum distance of 30 Å and confidence of 1. In addition, pocket-mode modelling was performed where every amino acid on the CDR3 loop of Nb23 was guided to an interaction with LRRK2 at a max distance of 6 Å and with confidence of 0.6. All the resulting models were filtered to remove redundancy and very low confidence hits.

### Statistical analysis

Data were checked for normal distribution using Shapiro-Wilk test. ROUT method to exclude outliers was performed. Normally distributed data were analyzed by One-way ANOVA or Two-way ANOVA, and Fisher’s LSD, Dunnett’s or Šídák’s multiple comparison. Non-normally distributed data were analyzed by Kruskal-Wallis test and Uncorrected Dunn’s multiple comparison. Significance values for all data are indicated in figure legends. Statistical analysis and graphs were generated using Prism software version 10.2.1 (GraphPad, USA).

## Results

### RAB29-mediated activation of LRRK2 is modified to varying degrees by LRRK2 phosphomutants

Previous studies have demonstrated that overexpression of RAB29 induces the recruitment of LRRK2 towards the Golgi, causing an hyperactivation of LRRK2 kinase activity with subsequent increases in pS1292 LRRK2 autophosphorylation and pT73 RAB10 substrate phosphorylation (Beilina et al., 2014; Liu et al., 2018; Madero-Pérez et al., 2018b; Purlyte et al., 2018). To determine the relevance of the phosphorylation status of LRRK2 in its activation, we tested the impact of RAB29-mediated activation on different phosphorylation mutants of LRRK2 by co-transfection of 3xFlag-tagged LRRK2 single and compound phosphomutants (6xS>A/D, S910A/D, S935A/D, S955A/D, S973A/D) and eGFP or eGFP-tagged RAB29 in HEK293T cells for 24h, and R1441G pathogenic LRRK2 mutant as active control (Figure 1). Co-expression of RAB29 and LRRK2 phosphomutants displayed a general increased autophosphorylation at S1292 LRRK2 and phosphorylation at T73 RAB10 with all the mutants when compared with LRRK2 phosphomutants expression alone (Figure 1C,D); and as expected, we did not observe a change of pS935 LRRK2 upon activation by RAB29 (Figure 1B). Phosphodead and phosphomimetic mutants activated by RAB29 displayed a similar increase of pS1292 LRRK2 and pT73 RAB10 when compared to WT LRRK2, with exception of single S910A and compound 6xS>A phosphodead LRRK2 mutants which caused a 2-3-fold higher increase of pS1292 LRRK2 and pT73 RAB10 relative to WT (Figure 1C,D).

**Figure 1.**
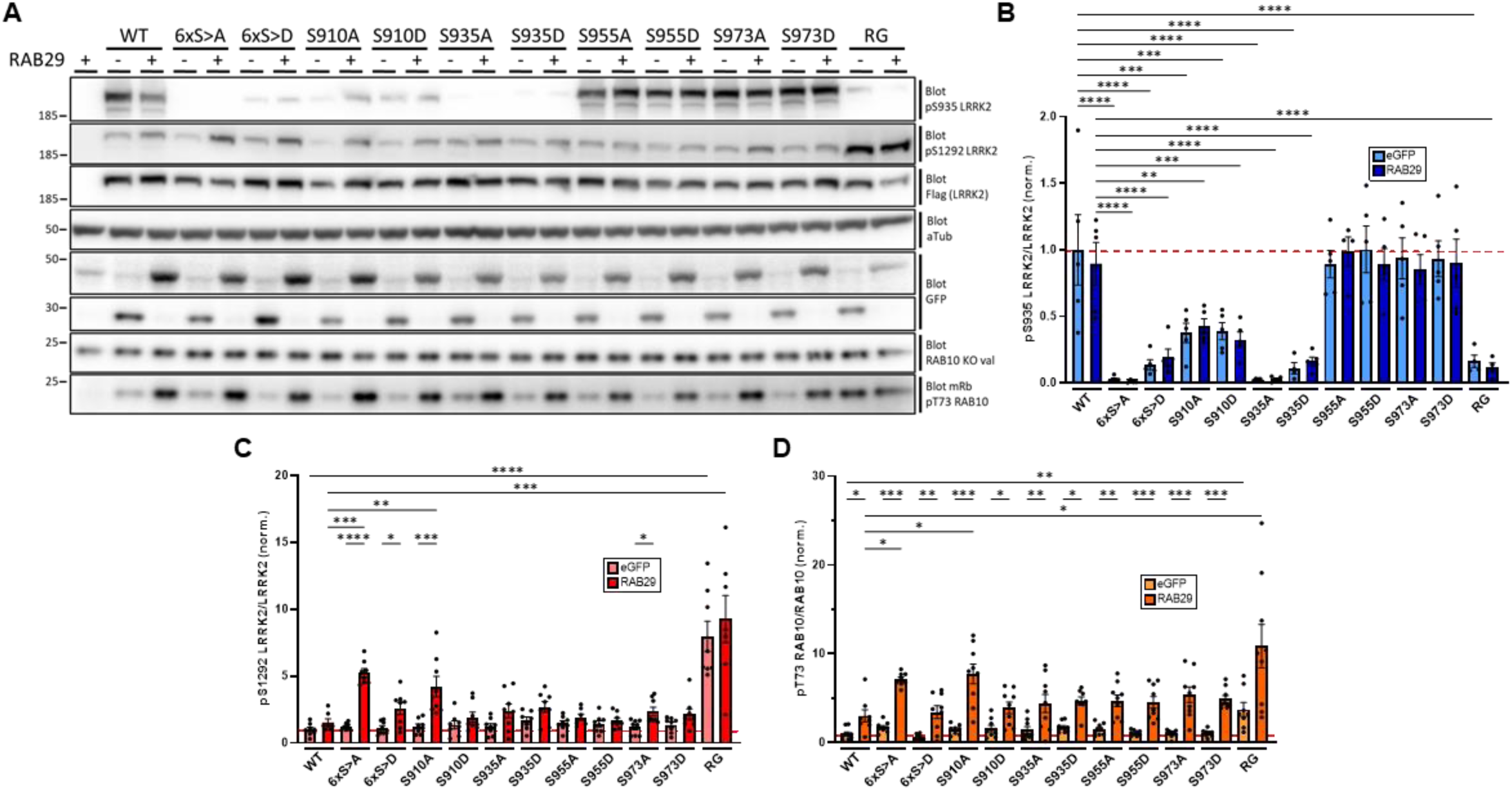
RAB29-mediated activation of LRRK2 is modified to varying degrees by LRRK2 phosphomutants. **(A)** HEK293T cells were transfected with the indicated 3xFlag-tagged LRRK2 construct and eGFP or eGFP-RAB29 where indicated, and extracts analyzed by western blotting for pS935 LRRK2, pS1292 LRRK2, Flag, GFP, pT73 RAB10, RAB10 and α-tubulin as loading control. **(B-D)** Quantification of pS935 LRRK2/LRRK2 ratio **(B)**, pS1292 LRRK2/LRRK2 ratio **(C)**, and pT73 RAB10/RAB10 ratio **(D)** of blots depicted in (A). Bars represent mean±s.e.m. (n=9 independent experiments); *p < 0.05, **p < 0.01, *** p < 0.005, **** p < 0.001.

### Time-course analysis of RAB29-mediated activation of LRRK2 shows that phosphodead LRRK2 is activated quicker and to a higher degree than phosphomimetic LRRK2

Co-expression of RAB29 and LRRK2 in HEK293T cells for 48h (Figure S2) displayed a milder activation, with lower LRRK2 kinase activity markers when compared with the expression at 24h (Figure 1). In order to determine whether there could be differences in activation over time, we performed a 48h time-course expression of RAB29 with WT LRRK2, 6xS>A/D and S910A/D mutants (Figure 2). Levels of phosphorylated S935 LRRK2 remained constant between the varying time points (Figure 2B). Interestingly, autophosphorylation of LRRK2 and RAB10 phosphorylation exhibited differential effects among phosphodead and phosphomimetic mutants (Figure 2C,D), suggesting a nuanced regulatory mechanism. We observed that LRRK2 autophosphorylation of phosphodead mutants by RAB29 overall reached its peak around 24 hours and was sustained over time. Conversely, the activation of phosphomimetic mutants by RAB29 exhibited a continuous increase during the evaluated time points (Figure 2C). In contrast, levels of LRRK2-mediated phosphorylation of RAB10 at residue T73 generally reached its peak of phosphorylation at 24h time point and, thereafter, it was reduced gradually for WT and all phosphomutants tested (Figure 2D). These observations were consistent with the findings from previous experiments conducted independently at 24 and 48 hours (Figure 1,S2).

**Figure 2.**
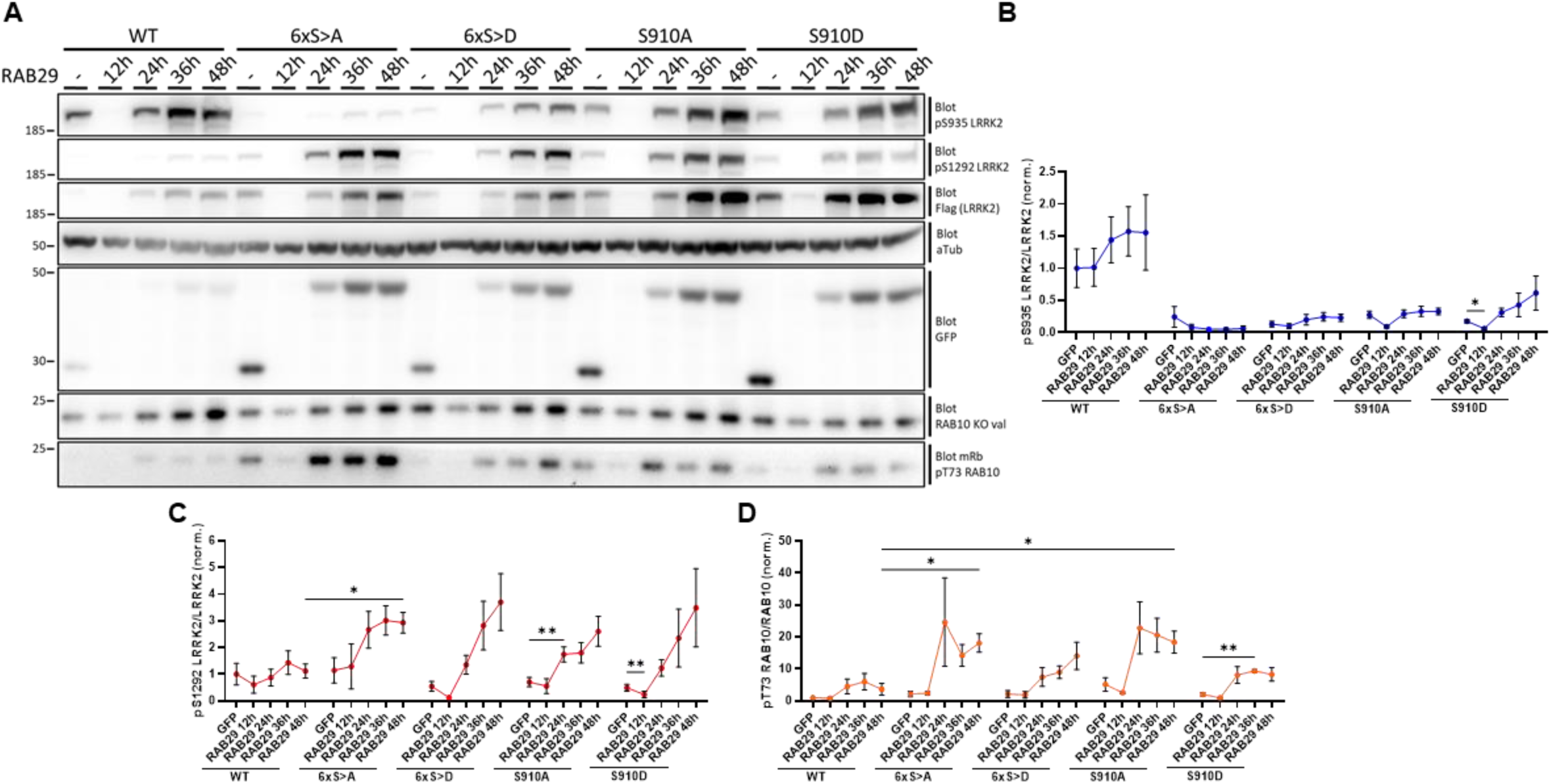
RAB29-mediated activation of LRRK2 depend on expression time. **(A)** HEK293T cells were transfected with the indicated 3xFlag-tagged LRRK2 construct and eGFP or eGFP-RAB29 where indicated, extracts collected at different time points and analyzed by western blotting for pS935 LRRK2, pS1292 LRRK2, Flag, GFP, pT73 RAB10, RAB10 and α-tubulin as loading control. **(B-D)** Quantification of pS935 LRRK2/LRRK2 ratio **(B)**, pS1292 LRRK2/LRRK2 ratio **(C)**, and pT73 RAB10/RAB10 ratio **(D)** of blots depicted in (A). Dots represent mean±s.e.m. (n=4 independent experiments); *p < 0.05, **p < 0.01.

### Analysis of RAB29-induced TGN recruitment of LRRK2 phosphomutants shows increased TGN recruitment for phosphodead LRRK2 compared to phosphomimetic LRRK2

As mentioned above, RAB29 expression has been reported to recruit LRRK2 towards TGN (Beilina et al., 2014; Liu et al., 2018; Madero-Pérez et al., 2018b; Purlyte et al., 2018). Therefore, we next asked whether this recruitment is affected by LRRK2 phosphorylation. We co-expressed 3xFlag-tagged LRRK2 single and compound phosphomutants in the presence or absence of eGFP-tagged RAB29 in HEK293T cells and analyzed the localization of LRRK2 phosphomutants respect to TGN by immunocytochemistry (Figure 3A). In all cases, LRRK2 phosphomutants were recruited towards TGN when co-expressed with RAB29 protein as assessed by analysis of co-localization of LRRK2 and RAB29 with a TGN marker (TGN46). Interestingly, we found that phosphomimetic mutants displayed a lower percentage of recruitment to Golgi when compared to its respective phosphodead mutant or WT LRRK2 (Figure 3B).

**Figure 3.**
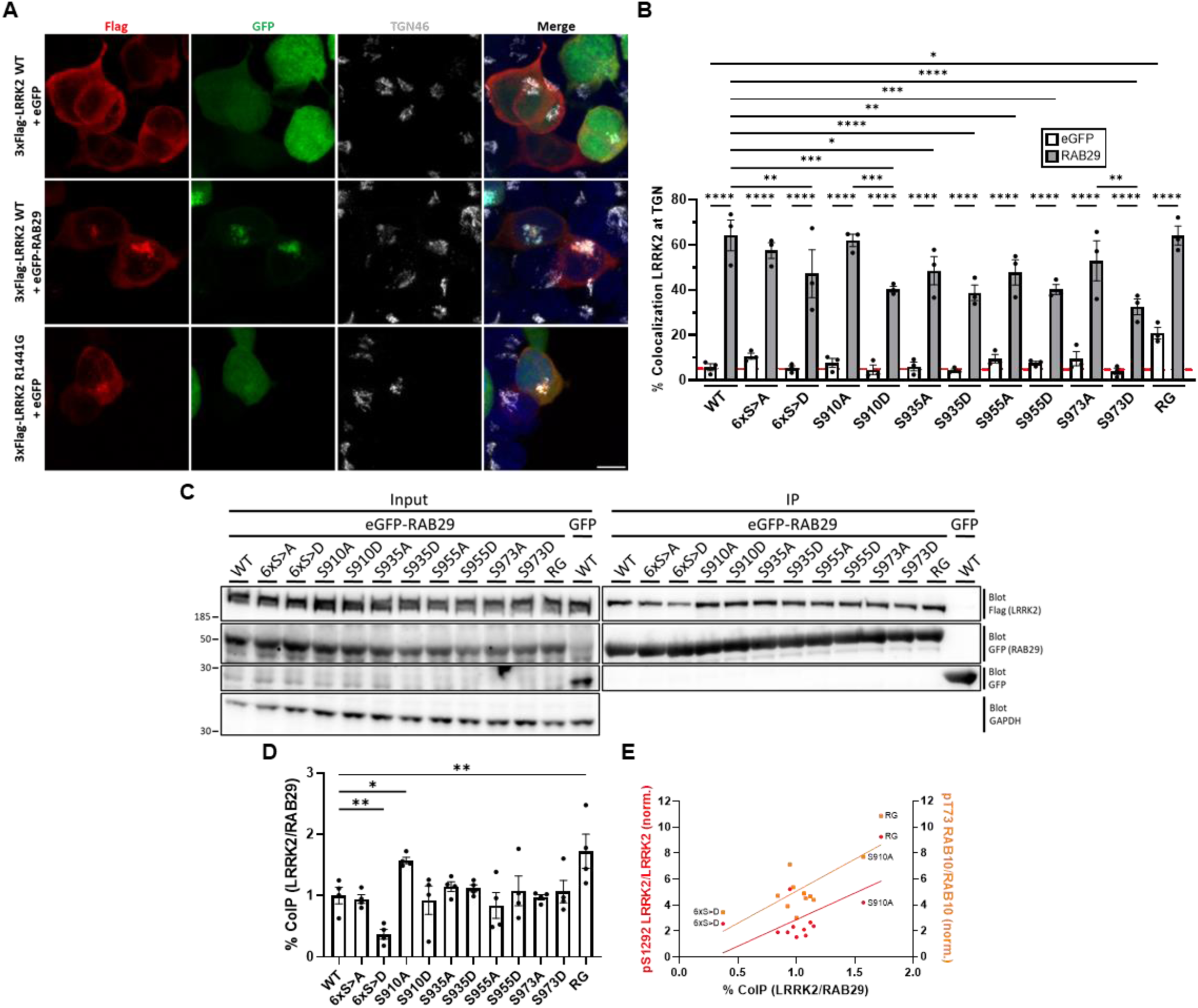
RAB29-mediated activation of LRRK2 depend on interaction and localization. **(A)** Example of HEK293T cells transfected with the indicated 3xFlag-tagged LRRK2 construct and eGFP or eGFP-RAB29 where indicated, and stained with antibodies against Flag (Alexa Fluor™ 568, red), Golgi marker TGN46 (Alexa Fluor™ 647, pseudocolored in gray) and DAPI. Scale bar, 10 µm. **(B)** Quantification of the percentage of cells co-transfected with LRRK2 and RAB29 that display colocalization of 3xFlag-tagged LRRK2 and eGFP-RAB29 at TGN from around 60 cells per condition. Bars represent mean±s.e.m. (n=3 independent experiments); *p < 0.05, **p < 0.01, *** p < 0.005, **** p < 0.001. **(C)** HEK293T cells were transfected with the indicated 3xFlag-tagged LRRK2 construct and eGFP or eGFP-RAB29 where indicated. Cell lysates were subjected for immunoprecipitation with anti-GFP magnetic beads and followed by western blot analysis with an anti-Flag or anti-GFP antibody. **(D)** Quantification of co-immunoprecipitation between LRRK2 and RAB29 of blots depicted in (C). Bars represent mean±s.e.m. (n=4 independent experiments);*p < 0.05, **p < 0.01. **(E)** Correlation plot of % coIP LRRK2/RAB29 and pS1292 LRRK2/LRRK2 ratio (red/left axis) and pT73 RAB10/RAB10 ratio (orange/right axis).

### Analysis of LRRK2:RAB29 interaction of LRRK2 phosphomutants shows increased interaction for phosphodead LRRK2 compared to phosphomimetic LRRK2

RAB29 binds LRRK2 in the ARM domain (Beilina et al., 2014; Vides et al., 2022; Zhu et al., 2023). We therefore next sought more direct evidence of a physical interaction between LRRK2 phosphomutants and RAB29 by co-immunoprecipitation (MacLeod et al., 2013). HEK293T cells were co-transfected with 3xFlag-tagged LRRK2 phosphomutants and eGFP-tagged RAB29 and, after 24h or 48h, cell lysates were immunoprecipitated with GFP-Trap® magnetic agarose and interaction assessed by western blot. GFP-immunoprecipitation of RAB29 effectively co-immunoprecipitated LRRK2 phosphomutants (Figure 3C,S3A). As observed for activation at different time points, we also observed some differences on interaction between LRRK2 and RAB29 depending on the time of expression. In general, interaction and co-immunoprecipitation between LRRK2 and RAB29 increased over time. At both times tested, R1441G pathogenic mutant displayed a significant increased interaction when compared to WT LRRK2 (Figure 3D,S3B). At 24 hours of transfection, phosphomimetic 6xS>D LRRK2 mutant displayed a significant reduced interaction with RAB29 when compared to WT LRRK2 (Figure 3D), whose difference became smaller upon expression for 48h (Figure S3B). According with the nuance differences observed for LRRK2 activation markers at 24h and 48h for S910 LRRK2 mutants, we also observed differences in the interaction between the phosphodead and the phosphomimetic mutants with RAB29. Phosphodead S910A LRRK2 mutant showed a significant increase interaction with RAB29 at 24h (Figure 3B), while phosphomimetic S910D LRRK2 mutant showed that increased interaction upon 48h of expression (Figure S3B). At both times tested, we found a positive correlation between the interaction of LRRK2:RAB29 and LRRK2 kinase activity markers (Figure 3E,S3C).

### A panel of LRRK2-targeting nanobodies stabilizing LRRK2 in varying states of phosphorylation show that LRRK2 stabilized at low S910/S935 phosphorylation is more sensitive to RAB29-mediated activation

A subset of nanobodies (Nbs) have been previously reported to bind LRRK2 at different domains and allosterically modulate LRRK2 activity (autophosphorylation and RAB phosphorylation) under RAB29-mediated activation conditions (Singh et al., 2022). To determine whether these Nbs could stabilize LRRK2 in a different heterologous phosphorylation state, we studied the effect of a panel of LRRK2-targeting Nbs on WT LRRK2. When 6xHis-GFP-tagged LRRK2-targeting nanobodies are co-expressed with LRRK2 in HEK293T cells (Figure 4A), we found that Nb22, Nb23 and Nb39 stabilize LRRK2 at a low S910 and S935 phosphorylation level, while this was not significantly altered by other Nbs (Figure 4B,C). In addition, in these basal conditions, Nb22 induced an activation of WT LRRK2 kinase activity displaying a significant increase of pS1292 LRRK2 and pT73 RAB10, while Nb23 caused an inhibition of LRRK2 kinase activity (Figure 4D,E).

**Figure 4.**
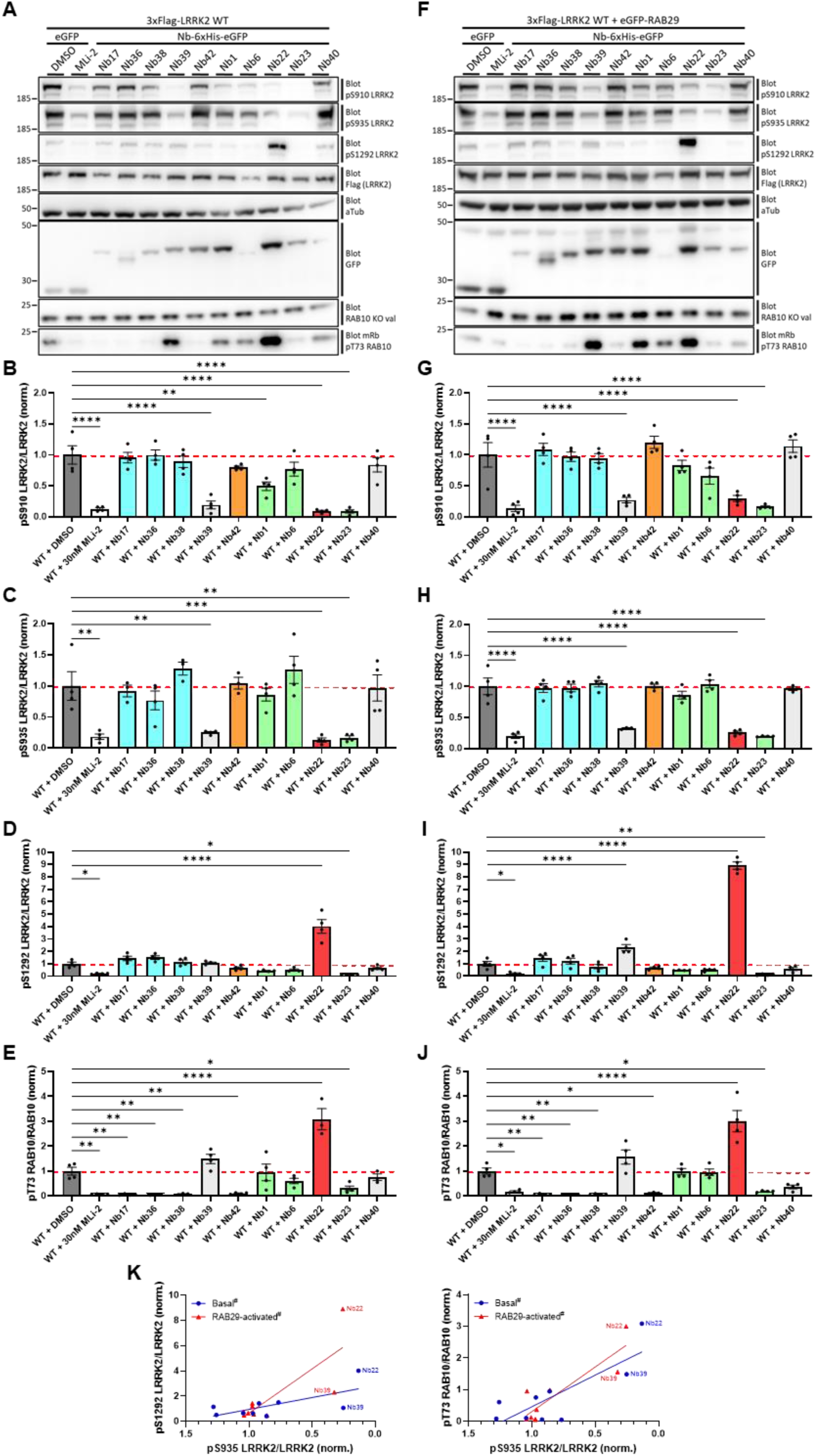
Impact of LRRK2-targeting nanobodies on LRRK2 kinase activity markers under basal or RAB29-mediated activation conditions. (A,F) HEK293T cells were transfected with 3xFlag-tagged WT LRRK2 with/without eGFP-RAB29, and eGFP or the indicated Nb-6xHis-eGFP where indicated, and extracts analyzed by western blotting for pS910 LRRK2, pS935 LRRK2, pS1292 LRRK2, Flag, GFP, pT73 RAB10, RAB10 and α-tubulin as loading control. **(B-E,G-J)** Quantification of pS910 LRRK2/LRRK2 ratio **(B,G)**, pS935 LRRK2/LRRK2 ratio **(C,H)**, pS1292 LRRK2/LRRK2 ratio **(D,I)**, and pT73 RAB10/RAB10 ratio **(E,H)** of blots depicted in (A,F). Bars represent mean±s.e.m. (n=4 independent experiments); *p < 0.05, **p < 0.01, *** p < 0.005, **** p < 0.001. **(K)** Correlation plot of pS935 LRRK2/LRRK2 ratio with pS1292 LRRK2/LRRK2 ratio (left) and pT73 RAB10/RAB10 ratio (right) under basal (blue) or RAB29-mediated activation (red) conditions. # Data points from the different conditions are normalized to their respective DMSO control.

We next characterized the effects of these Nbs on RAB29-activated WT LRRK2. When we co-expressed 3xflag-tagged LRRK2 and eGFP-tagged RAB29 in the presence of the different Nbs (Figure 4F), we observed the same reduction on LRRK2 heterologous phosphorylation (Figure 4G,H) when compared to basal LRRK2 with Nb22, Nb23 and Nb39 (Figure 4B,C). Under RAB29-activated conditions, LRRK2 also displayed an increased kinase activity when co-expressed with Nb22 and Nb39, and an inhibitory effect when co-expressed with Nb23 (Figure 4I,J), as similarly observed on RAB29-activated G2019S LRRK2. Interestingly, we found an inverse correlation between LRRK2 heterologous phosphorylation level and LRRK2 kinase activity in both basal and activation conditions which was more pronounced when kinase activity markers were tested in conditions of RAB29 activation, coherent with a higher sensitivity to RAB29 activation of dephosphorylated LRRK2 relative to basal LRRK2 (Figure 4K). The notable exception to this is the result for Nb23 that completely blocked LRRK2 kinase activity markers. Such a discrepancy could in principle be explained by a direct inhibition of substrate binding to LRRK2. Indeed, previous ELISA experiments showed that Nb23 binds to the kinase-WD40 region of LRRK2, while cross-linking mass spectrometry (XL-MS) experiments mapped it to the kinase domain (Singh et al., 2022). To obtain further insight into the binding mode of Nb23 we performed AlphaFold 3 (Abramson et al., 2024) and Chai-1 modeling (Chai Discovery, 2024), using either full-length LRRK2 or its kinase-WD40 domains indicating three potential binding positions for Nb23 (Figure S4-2A). Interestingly, when using previous XL-MS data as constraints, Nb23 docks in a position that would both interfere with the binding of substrates and is compatible with LRRK2 in its RAB29-activated conformation (Zhu et al., 2023) (Figure S4-2B). While further experimental validation of these predictions of the Nb23 binding position to LRRK2 will be required, we excluded Nb23 from the correlation analysis as it displays a profile similar to MLi-2, a type I kinase inhibitor of LRRK2 that both blocks kinase activity and also induces LRRK2 dephosphorylation at S910/S935.

The panel of nanobodies was tested also under RAB29 activation conditions on the phosphodead S910A LRRK2 mutant that we showed above to have increased sensitivity to RAB29 activation, as well as on the intrinsically hyperactive clinical G2019S LRRK2 mutant (Figure S4-1). We observed the same effects for Nb22, Nb23 and Nb39 on LRRK2 heterologous phosphorylation (Figure S4-1B,F,G) and LRRK2 kinase activity (Figure S4-1C,D,H,I) when compared to WT LRRK2. Interestingly for S910A LRRK2 mutant, in addition to being sensitized to RAB29 activation, it also inherently displays reduced pS935 LRRK2 levels (Figure 1B) and therefore correlation between LRRK2 heterologous phosphorylation and LRRK2 kinase activity markers in RAB29-activated conditions were flatter, as expected. On the other hand, the correlation was similar to WT for the G2019S LRRK2 pathogenic mutant (Figure S4-1J).

### Phosphodead 6xS>A LRRK2 mutant or activation of LRRK2 by RAB29 or Nb22 cause centrosome cohesion deficits

In order to test downstream cell physiological effects of LRRK2 at varying states of phosphorylation and activation, we verified whether centrosome cohesion deficits, a phenotype observed in cells expressing LRRK2 kinase activating clinical mutants and after RAB29 activation (Madero-Pérez et al., 2018a, 2018b; Fernández et al., 2019; Lara Ordóñez et al., 2019), could be modified in the presence of LRRK2 phosphomutants with and without RAB29 (Figure 5). Using the LRRK2 WT compared to both phosphodead and phosphomimetic hexamutants and S910 mutants with and without RAB29, we found as expected that all RAB29 conditions demonstrated increased centrosome cohesion deficits compared to conditions without RAB29 (Figure 5A). RAB29 conditions all showed about 50% cells with split centrosomes, corresponding to % observed also for the R1441G LRRK2 pathogenic mutant, suggesting that detrimental effects have reached a maximum in these conditions. Interestingly, 6xS>A and co-expression of WT LRRK2 and Nb22 conditions showed impaired centrosome cohesion without RAB29-mediated activation (Figure 5B). As both of these conditions displayed moderately enhanced kinase activity markers in basal conditions (i.e. without RAB29) (Figure 1,4), this suggests that the centrosome cohesion phenotype is rather sensitive to smaller changes in LRRK2 kinase activity with LRRK2 dephosphorylation.

**Figure 5.**
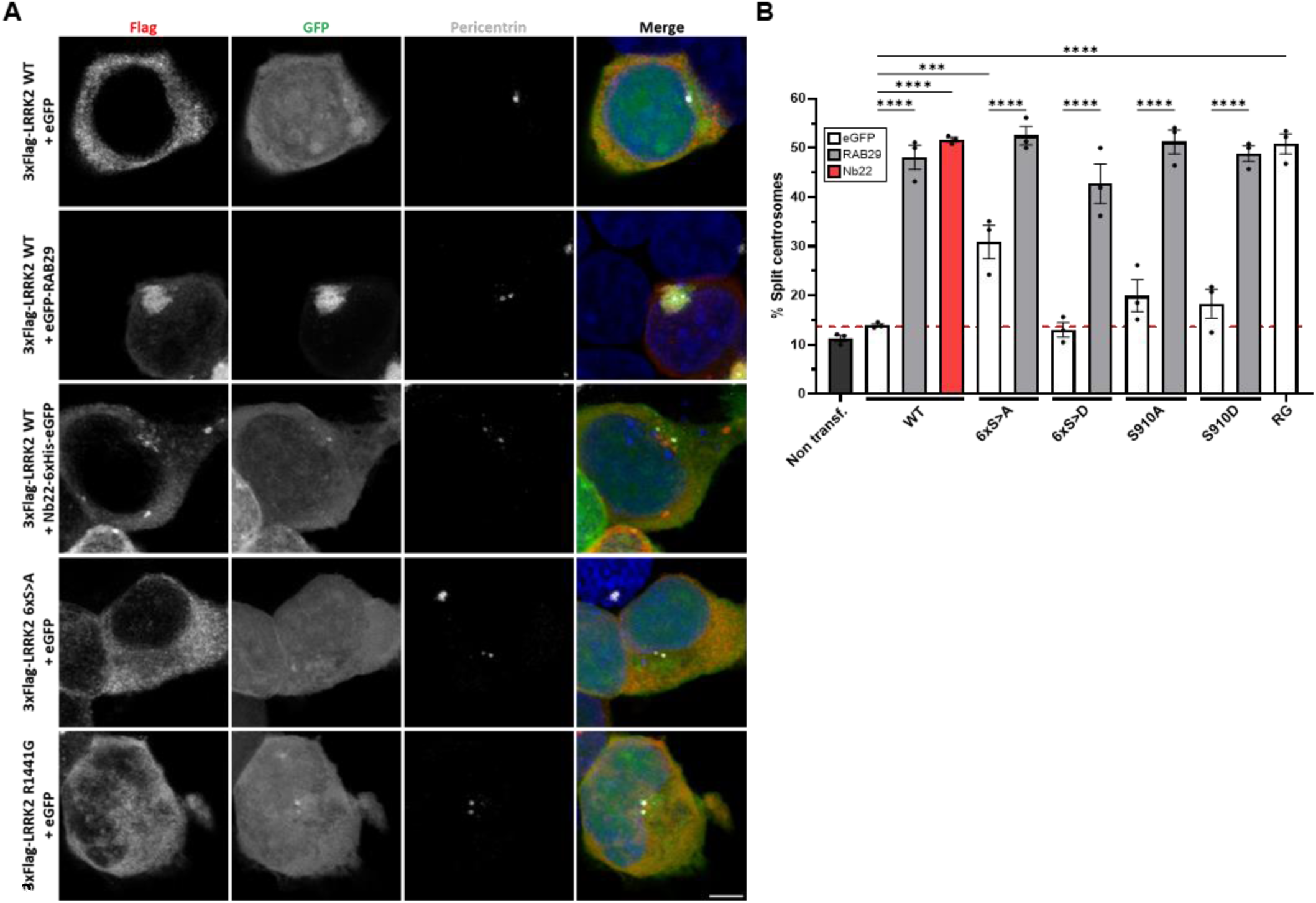
RAB29 and Nb22 activates LRRK2 phosphomutants causing centrosome cohesion deficits. **(A)** Example of HEK293T cells transfected with the indicated 3xFlag-tagged LRRK2 construct and eGFP, eGFP-RAB29 or Nb22-6xHis-eGFP where indicated, and stained with antibodies against Flag (Alexa Fluor™ 647, pseudocolored in red), pericentrin (Alexa Fluor™ 568, pseudocolored in gray) and DAPI. Scale bar, 5 µm. **(B)** Quantification of the percentage of split centrosome (duplicated centrosomes with a distance between their centers > 1.5 µm) from cells co-transfected with LRRK2 and eGFP, RAB29 or Nb22. Bars represent mean±s.e.m. (n=3 independent experiments); *** p < 0.005, **** p < 0.001.

## Discussion

In the present study, we tested how LRRK2 phosphorylation status at the heterophospholoop in the LRR domain impacts LRRK2 activation using both LRRK2 phosphomutants and nanobodies that stabilize LRRK2 in varying phosphorylation states. Our results show that hyperactivation of LRRK2 by upstream activator RAB29 is favored in conditions of low LRRK2 heterologous phosphorylation and disfavored in conditions of high LRRK2 heterologous phosphorylation.

Our study first confirmed (for the pS1292 LRRK2 autophosphorylation marker (Marchand et al., 2022) and extended (for the pT73-RAB10 substrate marker) previous findings that a compound phosphodead 6xS>A LRRK2 mutant showed close to 3-fold enhanced RAB29 activation compared to WT. When looking at the role of specific phosphosites in LRRK2 activation by RAB29, some sites stand out. For instance, upon RAB29 activation, the single phosphodead S910A LRRK2 mutant is hyperactivated to the same extent as the compound phosphodead 6xS>A LRRK2 mutant, suggesting that the S910 phosphosite may play a predominant role in RAB29-dependent activation of LRRK2. By contrast, the single phosphodead S935A LRRK2 mutant shows activation comparable to WT. Concerning the phosphomimetic mutants, we reproduce our previous findings that the compound phosphomimetic 6xS>D LRRK2 mutant displays reduced basal pT73 RAB10 rates (Marchand et al., 2022); however, upon activation with RAB29, we find that some single phosphomimetic mutants display activation apparently comparable to the corresponding single phosphodead mutant, particularly when observing the S1292 LRRK2 autophosphorylation. Interestingly, when we performed a time-course of the activation, we found a spike in the activation patterns of phosphodead mutants at 24h post-transfection that is not visible in the WT or phosphomimetic conditions, suggesting that dephosphorylated LRRK2 is subject to acute hyperactivation spikes upon RAB29 activation. By contrast, the phosphomimetic mutants, while retaining the ability to be activated by RAB29, do not display this spike in hyperactivation and we observe that the extent of activation and speed of activation is significantly lower for the phosphomimetic mutants, comparable to WT LRRK2.

While phosphomutants of LRRK2 offer insight into the role of individual sites compared to groups of sites in LRRK2 activation, there is the caveat that these constructs are approximations of true phosphorylation and dephosphorylation status. Therefore, we wanted to complement the phosphomutant studies with an additional approach that would allow us to relate LRRK2 phosphorylation state and sensitivity to activation. For this, we used a set of ten previously published LRRK2-targeting Nbs (Singh et al., 2022). An important property of Nbs is that they can promote specific conformations of the proteins they bind and we postulated that this set of Nbs would stabilize different conformations of LRRK2 and thereby drive different levels of LRRK2 heterologous phosphorylation. We co-expressed the Nbs fused to eGFP with LRRK2 and found a range of effects of the LRRK2-targeting Nbs on the S910 LRRK2 and S935 LRRK2 phosphorylation rates. When we tested LRRK2 kinase activity markers (pS1292 LRRK2 and pT73 RAB10) in the same basal conditions, we found an inverse correlation between LRRK2 heterologous phosphorylation and LRRK2 kinase activity. This is somewhat aligned with what is observed in the phosphomutants experiment, where some phosphodead mutants show a tendency to increased basal activity, although this did not reach statistical significance. We then tested the Nbs in conditions of RAB29 activation and found again a pronounced inverse correlation between LRRK2 heterologous phosphorylation and LRRK2 kinase activity. A side by side correlation plot of pS935 LRRK2 vs the two LRRK2 kinase activity markers for the Nbs with and without RAB29 show this similar trend (Figure 4K). In this plot, the RAB29-activated values are normalized to the control in RAB29 activation conditions, which has 3-fold enhanced activity compared to basal values without RAB29. Interestingly, for the pS1292 LRRK2 readout, the correlation plot is steeper suggesting synergistic effects of the dephosphorylation-inducing Nbs and RAB29 on activation for this readout. In the panel, there is a notable exception to this, namely Nb23 that shows low pS935 LRRK2 levels and also low autophosphorylation activity marker (pS1292 LRRK2). Although further structural validation is required, this exception could be explained by the binding of Nb23 on the kinase domain, where some of the modeled binding positions (Figure S4-2) suggest that Nb23 might interfere with RAB substrate binding, thereby directly blocking kinase activity. Such a mechanism of Nb23 would be somehow similar to what is observed with type I kinase inhibitors that also lead to dephosphorylation of S935 LRRK2 together with inhibition of LRRK2 kinase activity (Dzamko et al., 2010; Vancraenenbroeck et al., 2014; Fell et al., 2015). To complement these data on WT, we also tested two mutants in the presence of Nbs and RAB29, the S910A that shows increased sensitivity to RAB29 and the G2019S that has inherently higher kinase activity than WT. As expected, the correlation curves of pS935 LRRK2 vs kinase activity markers for the S910A LRRK2 are flatter, reflecting the lower pS935 LRRK2 values observed in phosphodead S910A LRRK2 mutant and a reduced contribution of phosphoregulating nanobodies in a variant that already shows greatly enhanced sensitivity to activation on its own. Interestingly, despite an inherent basal activation, the correlations between pS935 LRRK2 and kinase activity markers showed a similar trend for G2019S LRRK2 compared to WT, suggesting that phosphorylation levels are important for LRRK2 activation including for the G2019S LRRK2 mutant. Overall, these findings are consistent with our hypothesis that LRRK2 with low phosphorylation levels is more sensitive to activation than LRRK2 with higher phosphorylation levels, including for native LRRK2.

Besides testing how overexpression of activating RABs affects LRRK2 activation, other properties of activating RABs allow us to explore the activation mechanism in more detail, including binding of RAB29 to LRRK2 and RAB29-dependent recruitment of LRRK2 to TGN. Indeed, RAB29 is known to bind LRRK2 in the ARM domain and when we tested CoIP of RAB29 to the different phosphorylation mutants, we found that the extent of CoIP correlated to the degree of activation measured by pS1292 LRRK2 and pT73 RAB10 rates, suggesting that activation is linked to a higher degree of binding LRRK2:RAB29. We also tested recruitment by RAB29 of LRRK2 phosphomutants to TGN by immunocytochemistry. Here the effects were subtle with an overall trend showing stronger recruitment to TGN of phosphodead mutants compared to phosphomimetic mutants, which reached significance when comparing the recruitment of S910A vs S910D, and S973A vs S973D. Finally, we explored whether LRRK2 phosphorylation could affect centrosome cohesion deficits, a phenotype that has been linked to LRRK2 kinase activity and activation, including in patient derived lymphoblastoid cell lines. Indeed, centrosome cohesion deficits have been described as a robust readout that could be used to evaluate the efficacy of LRRK2 kinase inhibitors and as biomarker to stratify sporadic PD patients that could benefit from LRRK2-related therapies due to its sensitivity to detect increased LRRK2 kinase activity (Madero-Pérez et al., 2018a, 2018b; Fernández et al., 2019; Lara Ordóñez et al., 2019; Naaldijk et al., 2024). Here, we found that both the phosphodead 6xS>A LRRK2 mutant as well as a nanobody stabilizing dephosphorylated LRRK2 (Nb22) enhanced the centrosome cohesion deficit. Interestingly, all conditions with RAB29 activation, as well as the R1441G LRRK2 pathogenic mutant active control, showed a maximum effect of about 50% centrosome cohesion deficits. Also, the LRRK2-activating Nb22 showed a saturated effect in line with a 3-8-fold enhancement of LRRK2 activity in basal conditions. An intermediate effect is found for the phosphodead 6xS>A LRRK2 mutant that is shown to have about a 2-fold enhanced pT73 RAB10 marker level under basal conditions, corroborating that the centrosome cohesion deficit readout is very sensitive to LRRK2 activity.

It would be interesting to explore what the structural underpinnings are of the enhanced sensitivity to RAB29 activation of the S910 LRRK2 site, for instance whether this site is involved in direct or allosteric interaction with RAB29. At present, very little information is available on the fold and orientation of the heterophospholoop that is often unresolved in cryo-EM structures, likely due to the flexibility of the loop. RAB29 is described to land in the middle of the ARM domain (at armadillo repeats 9-10, positions 360-450, (Vides et al., 2022; Zhu et al., 2023), while the heterophospholoop is located in an extended loop of the LRR, positions 860-980. As we are showing increased activation of LRRK2 in dephosphorylated conditions, 14-3-3 is not bound to the S910/S935 sites, leaving the possibility that RAB29 may dynamically interact with the phospholoop, in particular the S910 residue, as part of the molecular events enhancing LRRK2 kinase hyperactivation. Further structural work will be required to determine how RAB29 and the LRRK2 heterophospholoop are coordinated during activation. It should be noted that the present study focuses on RAB29 and it remains important to verify whether LRRK2 heterologous phosphorylation status regulates sensitivity to activation but other upstream activators, including RAB12 (known to bind another region of the ARM domain), RAB32 or VPS35 that are both activators and PD risk factors.

The present findings have implications for biomarker interpretations and therapeutic targeting strategies. Indeed, levels of LRRK2 phosphorylation at S910/S935 are considered a biomarker of disease as lower levels are observed on average in disease compared to healthy controls, however interindividual variability makes that these measures are not sufficiently discriminating on their own (Dzamko et al., 2013; Fan et al., 2018; Fernández et al., 2019; Padmanabhan et al., 2020; Rideout et al., 2020). In light of the role these sites play in interaction, it may be of interest to look at the combination of the S910/S935 phospho-levels with expression levels of upstream activators, especially those that have been implicated in PD such as RAB29, but also RAB32, VPS35 or others. On therapeutic approaches, our study confirms that strategies that block LRRK2 dephosphorylation and promote LRRK2 phosphorylation are potentially viable therapeutic strategies, which offer an interesting alternative to approaches targeting LRRK2 catalytic sites or inhibit LRRK2 expression. Several targeting strategies would be possible based on our knowledge of LRRK2 phosphoregulation, including blocking access of phosphatases to the LRRK2 complex (Lobbestael et al., 2013; Athanasopoulos et al., 2016; Dong et al., 2020; Drouyer et al., 2021), favoring LRRK2 phosphorylation by upstream kinases or stabilization of the LRRK2:14-3-3 complex thereby favoring protection of LRRK2 phosphorylation sites. Interestingly, a cryo-EM structure of the LRRK2:14-3-3 complex, stabilized via crosslinking, shows LRRK2 in an inactive conformation and with 14-3-3 bound in a coordinated way above the S910 and S935 phosphorylation sites (Martinez Fiesco et al., 2025).

In conclusion, our study used a LRRK2-phosphomutant and LRRK2-nanobody approach to show that heterologous phosphorylation of LRRK2 determines the sensitivity of LRRK2 to activation by RAB29. In particular, our data points to phosphosite S910 as a driver of RAB29 activation of LRRK2. Mechanistically, LRRK2 phosphorylation dependent activation potential correlates to the degree of LRRK2:RAB29 binding and to RAB29-dependent recruitment of LRRK2 towards the TGN. Finally, measures of centrosome cohesion deficits link LRRK2 phosphorylation linked activity to a downstream pathological effect observed in PD. These findings suggest that the use of LRRK2 phosphorylation as a biomarker may be improved when combined with measures of LRRK2 upstream activators. Importantly, the results also validate an alternative targeting approach for LRRK2 based on promoting LRRK2 phosphorylation by targeting LRRK2 phosphoregulation complexes.

## Authors contribution

AJLO, CA and EB performed the experiments, data analysis, and statistical analysis; AJLO, CA, EB, MM, AB, LG and JMT contributed to data interpretation; FVM, CG and WV provided protein modelling analysis and material support; AJLO and JMT conceived and designed the study, and wrote the manuscript. All authors reviewed and approved the final manuscript.

## Acknowledgements

The authors are profoundly grateful for the funding from the Agence Nationale de la Recherche (ANR, grant ANR-21-CE16-0003-01 to JMT), the Michael J. Fox Foundation for Parkinson’s Research (MJFF, grants MJFF-022431, MJFF-025574, MJFF-023427 and MJFF-027170 to JMT), the Program Hauts-de-France FEDER-FSE+-FTJ 2021-2027 from the European Regional Development Fund (FEDER, grant HDF010441 to JMT and AJLO), the Strategic Research Program Financing from the VUB (grant SRP95 to WV) and the Research Foundation Flanders (grant G031324N to WV, and fellowship grant 1S01025N to FVM). We would like to thank Antonino Bongiovanni and Sophie Desnoulez from the Photonic Microscopy Facility of BioImaging Center Lille (UAR 2014 - US 41 - PLBS, F-59000 Lille, France) for the expert technical assistance.

---

**Figure S2:**
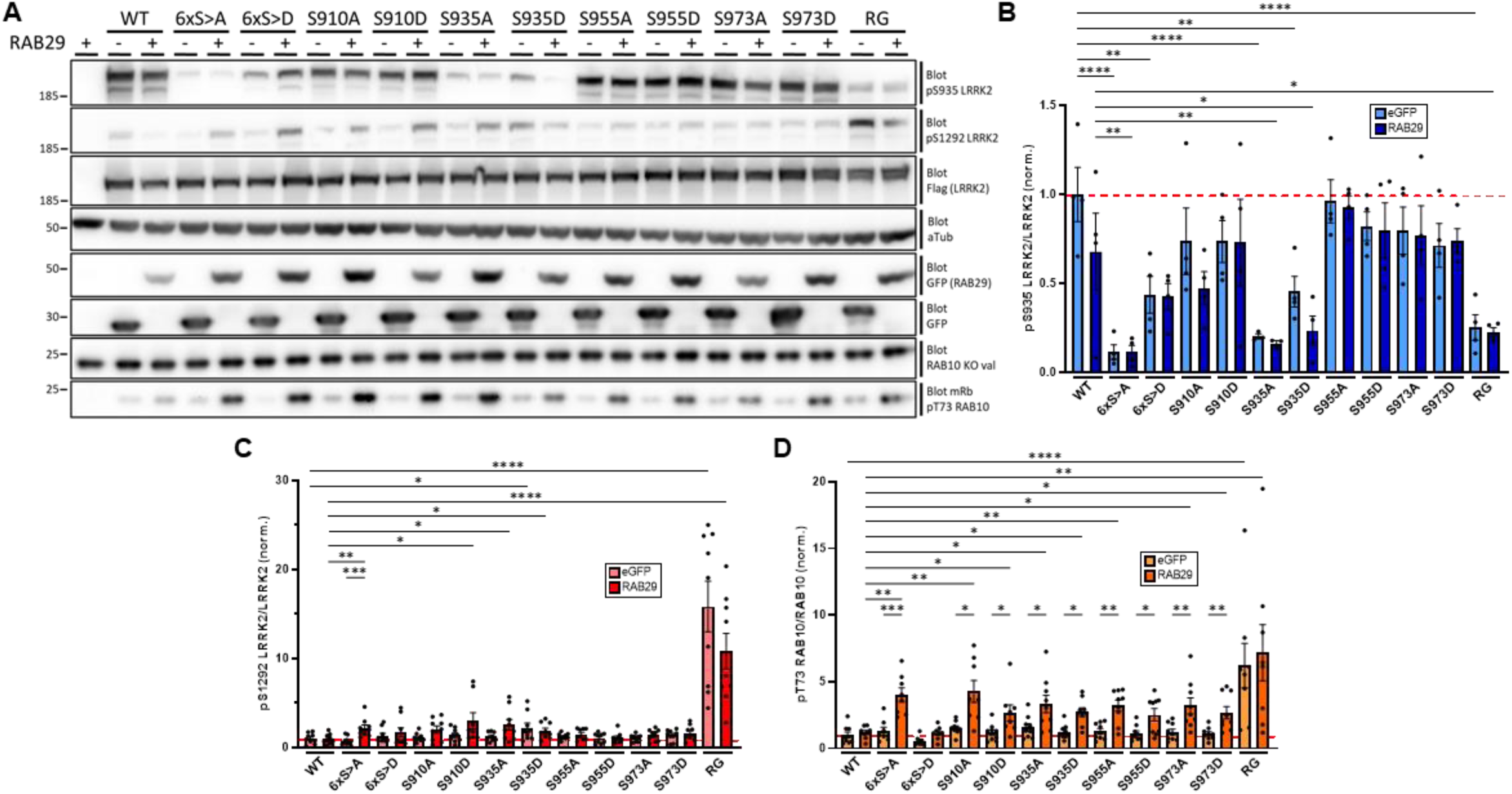
RAB29-mediated activation of LRRK2 phosphomutants upon 48h of expression. **(A)** HEK293T cells were transfected with the indicated 3xFlag-tagged LRRK2 construct and eGFP or eGFP-RAB29 where indicated, and extracts analyzed by western blotting for pS935 LRRK2, pS1292 LRRK2, Flag, GFP, pT73 RAB10, RAB10 and α-tubulin as loading control. **(B-D)** Quantification of pS935 LRRK2/LRRK2 ratio **(B)**, pS1292 LRRK2/LRRK2 ratio **(C)**, and pT73 RAB10/RAB10 ratio **(D)** of blots depicted in (A). Bars represent mean±s.e.m. (n=9 independent experiments); *p < 0.05, **p < 0.01, *** p < 0.005, **** p < 0.001.

**Figure S3.**
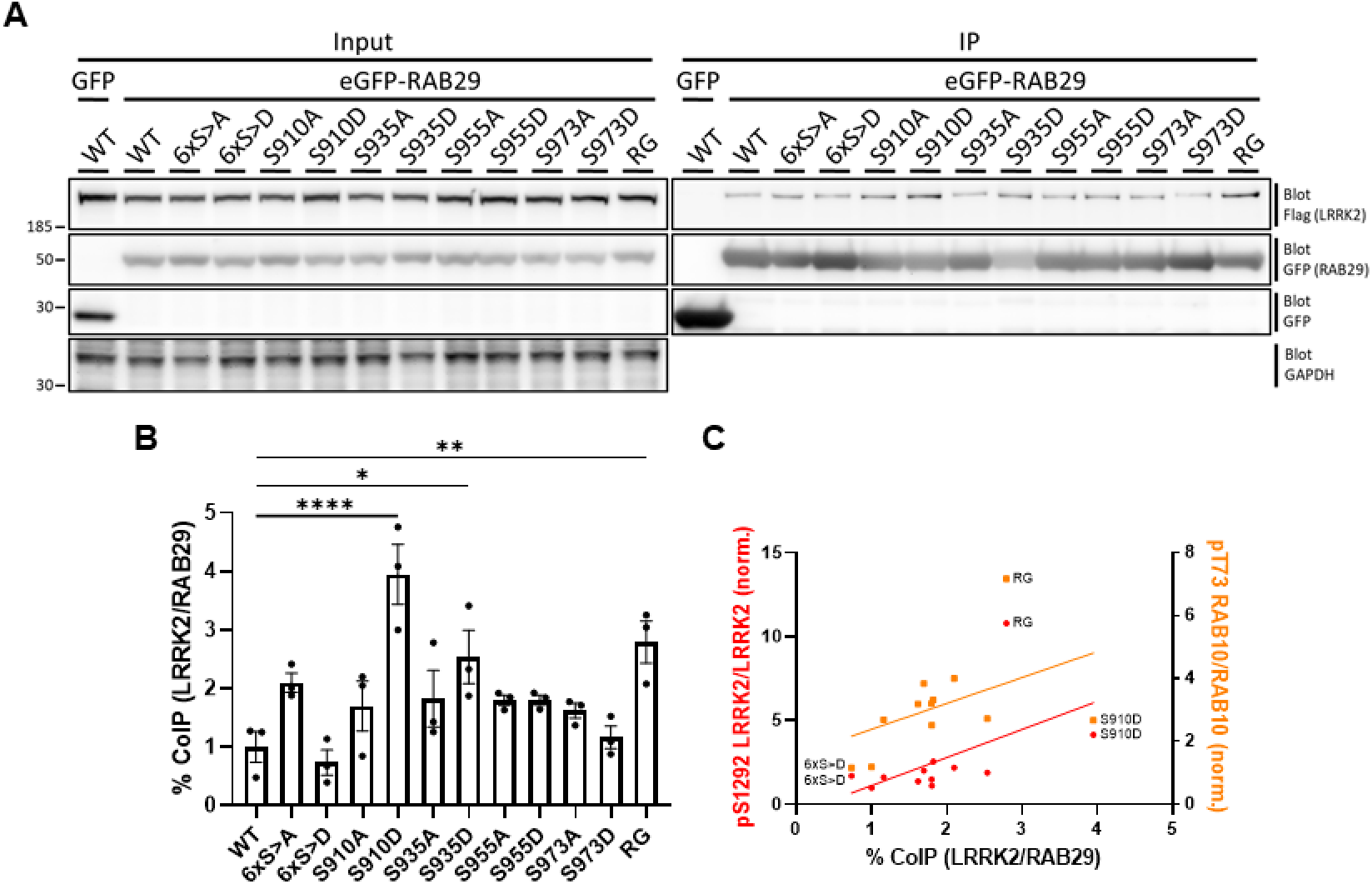
Co-immunoprecipitation of LRRK2 and RAB29 upon 48h of expression. **(A)** HEK293T cells were transfected with the indicated 3xFlag-tagged LRRK2 construct and eGFP or eGFP-RAB29 where indicated. Cell lysates were subjected for immunoprecipitation with anti-GFP magnetic beads and followed by western blot analysis with an anti-Flag or anti-GFP antibody. **(B)** Quantification of co-immunoprecipitation between LRRK2 and RAB29 of blots depicted in (A). Bars represent mean±s.e.m. (n=4 independent experiments);*p < 0.05, **p < 0.01, ****p < 0.001. **(C)** Correlation plot of % coIP LRRK2/RAB29 and pS1292 LRRK2/LRRK2 ratio (red/left axis) and pT73 RAB10/RAB10 ratio (orange/right axis).

**Figure S4-1.**
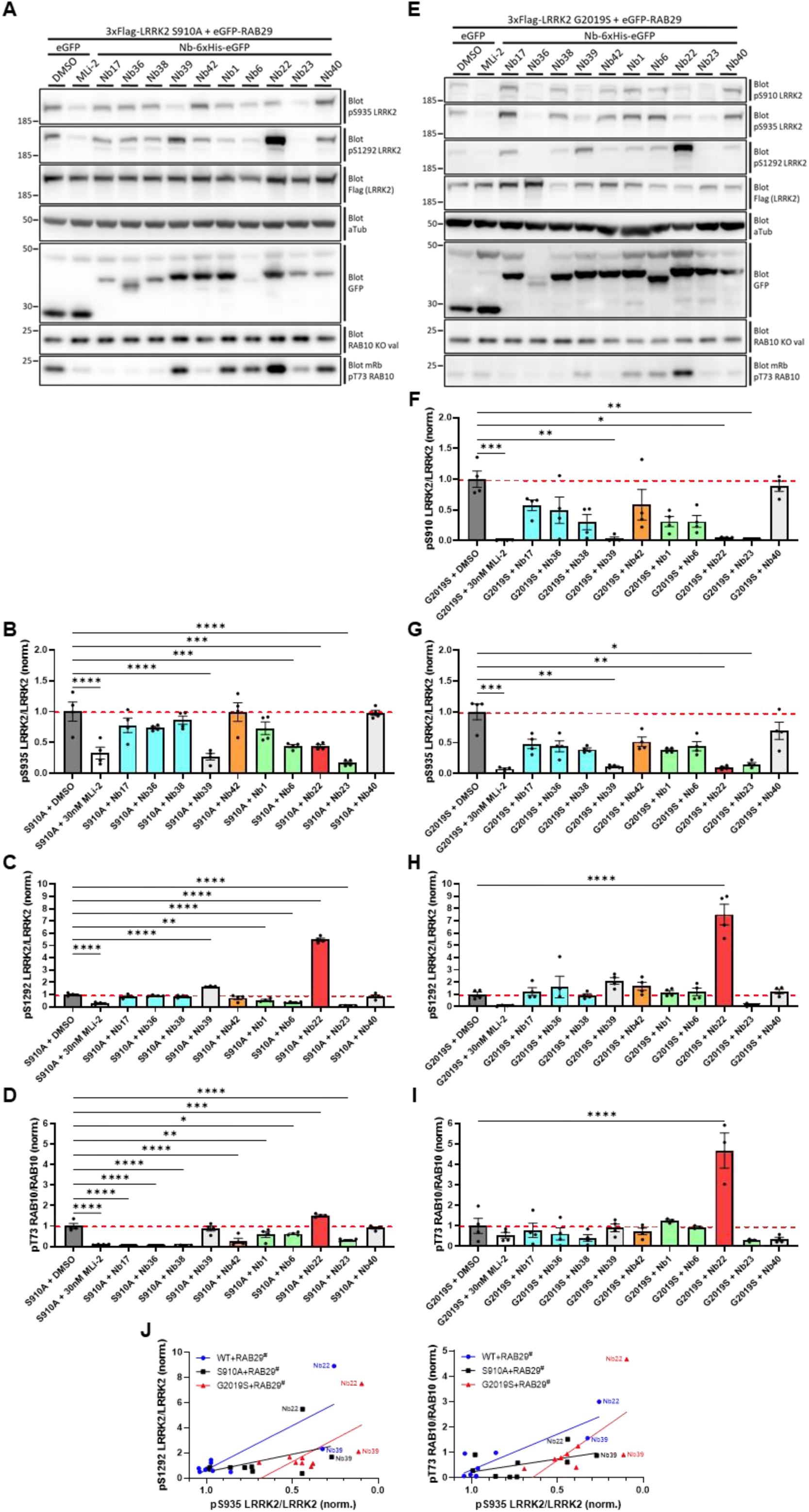
Impact of LRRK2-targeting nanobodies on LRRK2 kinase activity markers of S910A and G2019S mutants under RAB29-mediated activation conditions. (A,E) HEK293T cells were transfected with the indicated 3xFlag-tagged LRRK2 and eGFP-RAB29, and eGFP or the indicated Nb-6xHis-eGFP where indicated, and extracts analyzed by western blotting for pS910 LRRK2, pS935 LRRK2, pS1292 LRRK2, Flag, GFP, pT73 RAB10, RAB10 and α-tubulin as loading control. **(B-D,F-I)** Quantification of pS910 LRRK2/LRRK2 ratio **(F)**, pS935 LRRK2/LRRK2 ratio **(B,G)**, pS1292 LRRK2/LRRK2 ratio **(C,H)**, and pT73 RAB10/RAB10 ratio **(D,I)** of blots depicted in (A,E). Bars represent mean±s.e.m. (n=4 independent experiments); *p < 0.05, **p < 0.01, *** p < 0.005, **** p < 0.001. **(J)** Correlation plot of pS935 LRRK2/LRRK2 ratio with pS1292 LRRK2/LRRK2 ratio (left) and pT73 RAB10/RAB10 ratio (right) for RAB29-mediated activation of WT LRRK2 (blue), S910A LRRK2 mutant (black) and G2019S LRRK2 pathogenic mutant (red). # Data points from the different conditions are normalized to their respective DMSO control.

**Figure S4-2.**
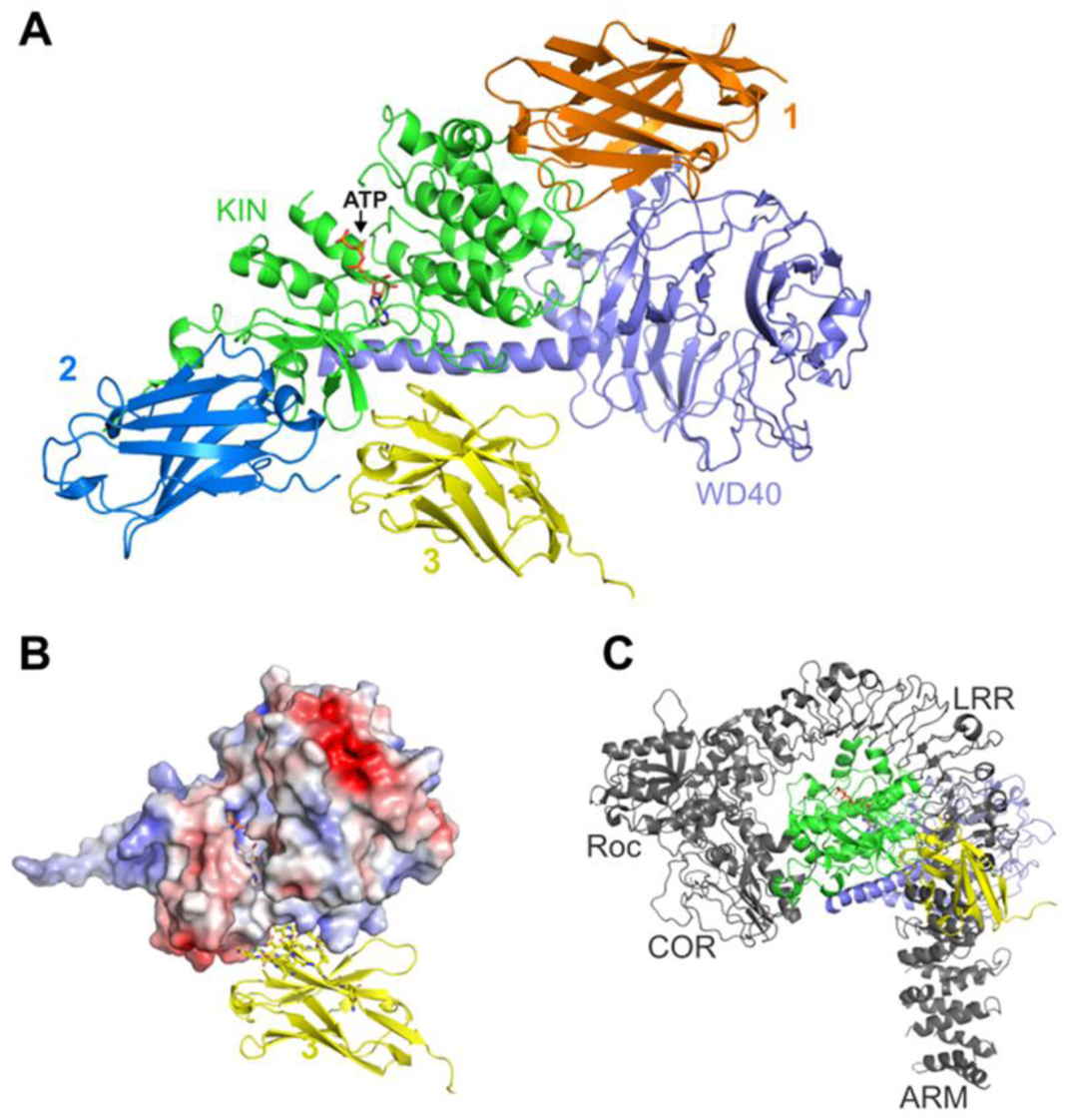
AlphaFold 3 (AF3) and Chai-1 modeling of potential binding modes of Nb23 to the LRRK2 kinase-WD40 domains. **(A)** Three predicted binding positions of Nb23, with Site 1 (Nb23 in orange) predicted by both AF3 and Chai-1 that docks Nb23 on the kinase domain in a region spatially close to the WD40 domain, Site 2 (Nb23 in blue) predicted by AF3 places Nb23 on the kinase domain in a position close to the connection with the COR domain, and Site 3 (Nb23 in yellow) predicted by Chai-1 using previous XL-MS data as constraints (Singh et al., 2022) places Nb23 closer to the ATP binding pocket of the kinase domain. The LRRK2 kinase-WD40 domains are shown with the kinase in green and the WD40 in light blue. **(B)** Zoom-in on the “Site 3” binding mode of Nb23. The kinase domain is shown in electrostatic potential surface representation. The ATP molecule in the active site is shown in stick representation. **(C)** Superposition on full-length LRRK2 shows that Nb23 in Site 3 would sterically clash with the LRRK2 N-terminal domains when LRRK2 adopts its compact auto-inhibited state.

